# Pleiotropy of autism-associated chromatin regulators

**DOI:** 10.1101/2022.12.07.519375

**Authors:** Micaela Lasser, Nawei Sun, Yuxiao Xu, Karen Law, Silvano Gonzalez, Belinda Wang, Vanessa Drury, Sam Drake, Yefim Zaltsman, Jeanselle Dea, Ethel Bader, Kate E. McCluskey, Matthew W. State, A. Jeremy Willsey, Helen Rankin Willsey

## Abstract

Gene ontology analyses of high confidence autism spectrum disorder (hcASD) risk genes have historically highlighted chromatin regulation and synaptic function as major contributors to pathobiology. Our recent functional work *in vivo* has additionally implicated microtubule biology and identified disrupted cellular proliferation as a convergent ASD phenotype. As many chromatin regulators, including ASD risk genes *ADNP* and *CHD3*, are known to directly regulate both tubulins and histones, we studied the five chromatin regulators most strongly associated with ASD (*ADNP, CHD8, CHD2, POGZ*, and *SUV420H1/KMT5B*) specifically with respect to microtubule biology. We observe that all five localize to microtubules of the mitotic spindle *in vitro* and *in vivo*. Further in-depth investigation of *CHD2* provides evidence that patient-derived mutations lead to a range of microtubule-related phenotypes, including disrupted localization of the protein at the mitotic spindle, spindle defects, cell cycle stalling, DNA damage, and cell death. Lastly, we observe that ASD genetic risk is significantly enriched among microtubule-associated proteins, suggesting broader relevance. Together, these results provide further evidence that the role of tubulin biology and cellular proliferation in ASD warrant further investigation and highlight the pitfalls of relying solely on annotated gene functions in the search for pathological mechanisms.

## INTRODUCTION

The past two decades have yielded striking progress in the identification of high-confidence (hc), large-effect risk genes for autism spectrum disorders (ASD), with robust statistical support for over 100 hcASD genes (Fu et al., 2022; Willsey et al., 2022). Gene ontology analyses of risk genes have repeatedly emphasized enrichment of terms related to synaptic biology and gene expression regulation by transcription factors and chromatin regulators (De Rubeis et al., 2014; Iossifov et al., 2012; Neale et al., 2012; Sanders et al., 2012; Satterstrom et al., 2020). At the same time, our hypothesis-naive *in vivo* investigation of the ten genes with the strongest statistical evidence for association with ASD has identified cell proliferation as an additional point of functional convergence (Willsey et al., 2021), consistent with other work in patient-derived induced pluripotent stem cells (iPSCs) (Adhya et al., 2021; Marchetto et al., 2017; Mariani et al., 2015). In orthologous work, focusing on the hcASD genes *DYRK1A* and *KATNAL2*, we identified evidence for the involvement of microtubule dysfunction (Willsey et al., 2018, 2020), raising the possibility that disruption of the mitotic spindle, via alteration of tubulin biology, may explain, at least in part, the proliferation phenotypes among hcASD gene mutants. Consistent with this possibility, many chromatin regulators, including the ASD-associated genes *ADNP* and *CHD3*, are known to modulate both histones and tubulins (Bassan et al., 1999; Koenning et al., 2021; Mandel and Gozes, 2007; Mandel et al., 2007; Matsuoka et al., 2008; Ostapcuk et al., 2018; Oz et al., 2014; Park et al., 2016; Sillibourne et al., 2007; Yokoyama et al., 2013). These findings suggest that ASD-associated chromatin regulators may play pleiotropic roles in general, regulating both histones and tubulins and that disruptions of one, the other, or both processes may underlie ASD pathobiology. However, this has been unexplored.

Thus, we sought to test two hypotheses: first, that ASD-associated genes annotated as chromatin regulators have dual functions in regulating chromatin and tubulin and second, that alterations in microtubule-related processes are related to ASD pathobiology in general. To address the first question, we turned to the *in vivo Xenopus tropicalis* model system as well as *in vitro* human tissue culture model systems. We observe that the five chromatin regulators most strongly associated with ASD (ADNP, CHD8, CHD2, POGZ, and SUV420H1/KMT5B) (Fu et al., 2022) do indeed localize to microtubules *in vivo* and *in vitro*. Focusing on *CHD2*, we demonstrate that recapitulating ASD-associated haploinsufficiency *in vitro* results in phenotypes consistent with a role in regulating microtubules, including mitotic spindle defects, cell cycle progression defects, DNA damage, and cell death. Moreover, we demonstrate that an ASD patient-derived missense mutation disrupts the localization of CHD2 to the mitotic spindle *in vivo*. To address our second hypothesis, namely that derangements in microtubule function contribute to ASD pathobiology *writ large*, we leveraged the past decade of successful gene discovery in ASD in conjunction with experimentally-derived tubulin-centric protein-protein interaction (PPI) data. We determine that hcASD genes are overrepresented among tubulin-related proteins and that ASD patient-derived protein truncating variants are more likely to occur within tubulin-related proteins. Overall, these findings strongly suggest that disruption of microtubule biology is an underappreciated component of ASD pathobiology, underscoring the limitations of relying on annotated function to identify convergent pathological mechanisms in ASD, and highlighting the importance of hypothesis-free functional exploration of risk genes to help clarify pleiotropy. Further, as chromatin regulators are implicated in risk for a large variety of psychiatric disorders, congenital heart disease, and cancer, these results may similarly suggest a broader role for microtubule biology in human disease.

## RESULTS

### ASD-associated chromatin regulators localize to microtubules

To test our hypothesis that ASD-associated chromatin regulators may also regulate tubulins, we visualized the subcellular localizations of the five chromatin regulators with the strongest statistical evidence for association with ASD (ADNP, CHD8, CHD2, POGZ, and SUV420H1/KMT5B) (Fu et al., 2022). First, we expressed strep-tagged human cDNAs for these genes *in vivo* in *Xenopus* embryos and imaged interphase and mitotic cells, staining for the strep tag and labeling DNA with DAPI. In interphase cells, ASD-associated chromatin regulators ADNP, CHD8, CHD2, POGZ, and SUV420H1/KMT5B localized to the nucleus, consistent with their reported functions as gene expression regulators (**Fig. 1A**). However, in metaphase cells, all five proteins localized to microtubule-rich mitotic spindles (**Fig. 1B**). We confirmed these localizations *in vitro* in mitotic human HEK293 cells (**Fig. S1**) and negative control strep experiments did not show background staining *in vivo* or *in vitro* (**Fig. 1, S1**). Together, these data indicate that the five most strongly ASD-associated chromatin regulators localize to microtubules in multiple cell types across both *in vivo* and *in vitro* models, consistent with potential functional roles of these proteins in regulating microtubules.

**Figure 1.**
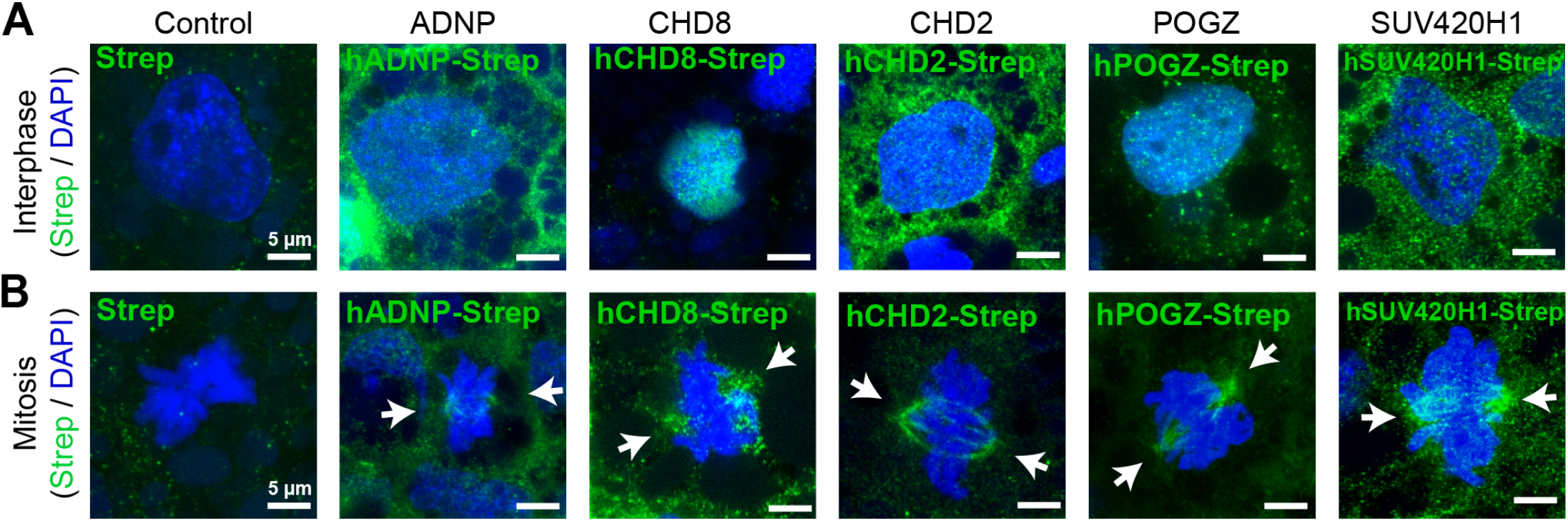
ASD-associated chromatin regulators localize to microtubules *in vivo*. (A) Human strep-tagged constructs for ASD-associated chromatin regulators ADNP, CHD8, CHD2, POGZ, and SUV420H1/KMT5B (labeled by Strep, green) localize to the nucleus (labeled by DAPI, blue) during interphase when expressed in *Xenopus*. (B) However, these constructs localize to the mitotic spindle during mitosis. Negative control uninjected animals stained with the strep antibody do not show these localizations. ADNP is considered a positive control since it has reported functions in gene expression regulation and microtubule stability. See Figure S1 for localizations in HEK293 cells, which support these observations.

### CHD2 is required for spindle organization, cell cycle progression, genome stability, and survival

Given this evidence that multiple, high-confidence ASD risk genes involved in chromatin biology encode proteins that also localize to the mitotic spindle, we next sought to interrogate whether ASD-associated mutations alter microtubule biology. As many large-effect mutations contributing to ASD lead to haploinsufficiency, we characterized the impact of *CHD2* loss-of-function on mitosis, a proliferation-related process that depends on dynamic microtubule regulation. To do this, we inhibited *CHD2* expression via CRISPRi in human induced pluripotent stem cell (iPSC) derived cortical neural progenitor cells (NPCs), an *in vitro* model of a cell type we and others have previously implicated in ASD risk (Willsey et al., 2021, 2022). We confirmed *CHD2* knockdown efficiency by qPCR (**Fig. S2A**) and assayed whether mitotic spindle organization was disrupted. We observed that reduction of *CHD2* led to an increase in the frequency of abnormal mitotic spindles (labeled by β-tubulin) in dividing cells (labeled by pHH3 staining) compared to non-targeting CRISPRi controls (**Fig. 2A, 2E, S2B**; Chi-squared test p = 0.047). Defects in spindle organization often lead to cell cycle stalling, DNA damage, and cell death; therefore, we assessed each of these phenotypes. We assayed cell cycle progression by staining NPCs with cell cycle checkpoint markers, Cyclin B for G2/M phases and Cyclin E for G1/S phases, and quantifying fluorescence intensity across cells. We observed that *CHD2* reduction caused an increase in Cyclin B (**Fig. 2B, F**; Rank sum test p < 0.0001) and a decrease in Cyclin E compared to non-targeting CRISPRi controls (**Fig. S2C-D**; Rank sum test p < 0.0001), suggesting that a larger proportion of NPCs were in G2/M phases. This is the expected result if the mitotic spindle checkpoint is activated due to abnormal spindle organization and mis-segregation of chromosomes (Bharadwaj and Yu, 2004). This would lead to cell cycle stalling during M phase, causing an increase in the proportion of cells positive for Cyclin B (G2/M marker) and a decrease in the proportion of cells in other cell cycle phases (Cyclin E, G1/S marker). Abnormal spindle organization also often causes DNA damage, as the spindle cannot properly segregate chromosomes. We therefore sought to determine whether DNA damage was increased following reduction of *CHD2* expression. To do this, we used an antibody against phosphorylated H2A histone family member X (pH2AX), a canonical marker for improper cell division and the resulting chromosomal instability (Rogakou et al., 1998), and quantified the number of pH2AX puncta per nucleus. We observed that loss of *CHD2* significantly increased the number of pH2AX puncta per nucleus compared to controls (**Fig. 2C, G**; Rank sum test p < 0.0001). This result is consistent with a study observing a higher rate of DNA damage in iPSC-derived NPCs from patients with ASD (Wang et al., 2020). DNA damage can trigger apoptosis, and we also observed an increase in cell death, as measured by the proportion of cells positive for cleaved caspase 3 antibody staining (**Fig. 2D, H**; Rank sum test p < 0.0001). Together, these data point to a role for CHD2 in spindle organization, cell cycle progression, genome stability, and cell survival in human NPCs, which could be due to a function in regulating microtubule dynamics at the mitotic spindle.

**Figure 2.**
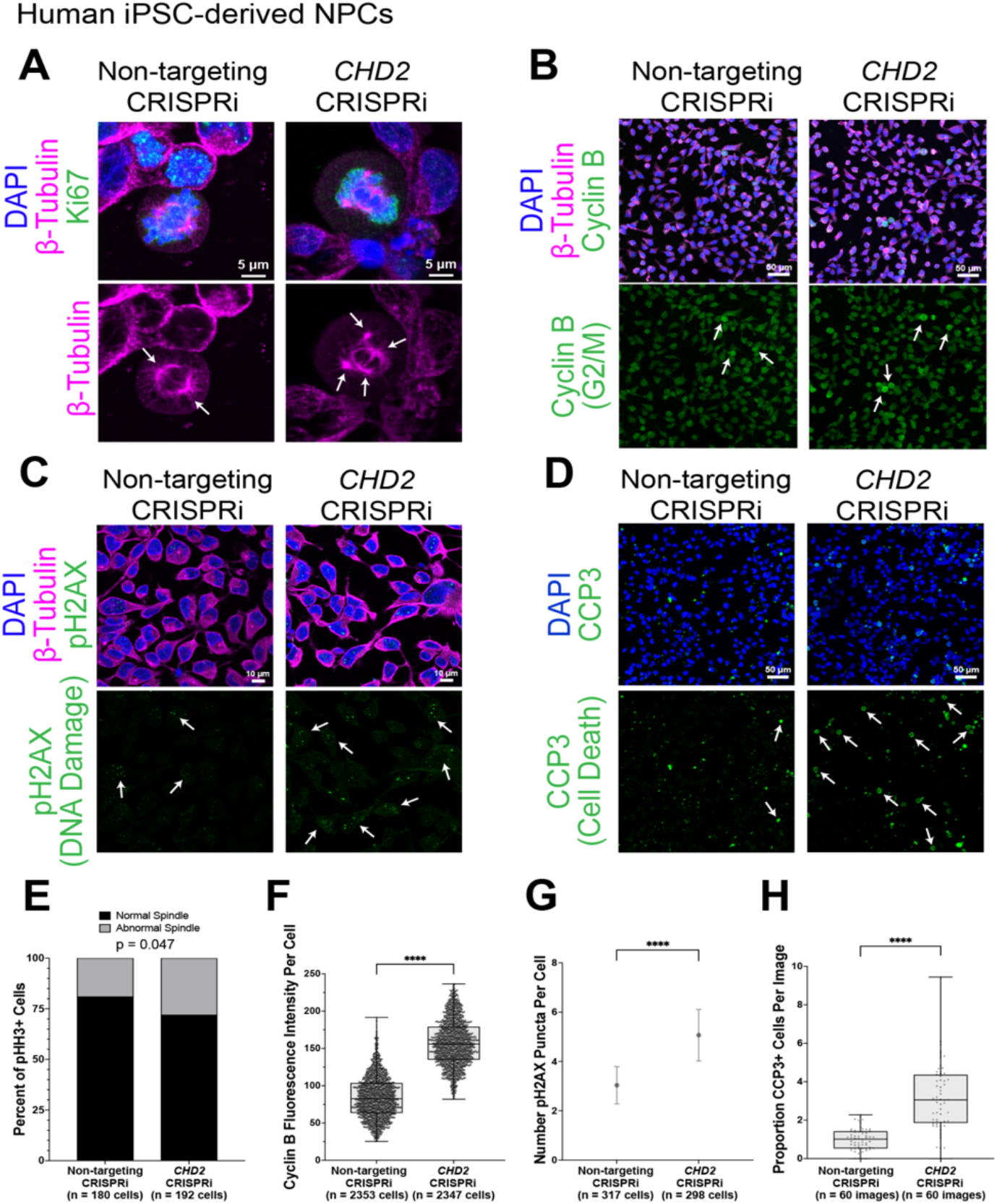
CHD2 is required for mitotic spindle organization, cell cycle progression, genome stability, and survival. (A) CRISPRi of *CHD2* in human iPSC-derived neural progenitor cells causes an increase in mitotic spindle defects (arrows, multipolar spindle) compared to non-targeting sgRNA controls (left, bipolar spindle). (B) CRISPRi of *CHD2* causes an increase in Cyclin B (G2/M marker) fluorescence per cell compared to a non-targeting CRISPRi line. (C) *CHD2* CRISPRi causes an increase in the number of phospho-Histone H2AX (pH2AX, DNA damage marker) puncta per nucleus compared to non-targeting CRISPRi. (D). *CHD2* CRISPRi causes an increase in the proportion of cleaved caspase 3 (CCP3, cell death marker) positive cells compared to non-targeting CRISPRi. (E) Quantification of (A). Chi squared test. (F) Quantification of (B). Box is 25-75% interquartile range, line is median, and whiskers are max to min. (G) Quantification of (C). Dot is at the mean and lines are 95% confidence intervals. (H) Quantification of (D). Box is 25-75% interquartile range, line is median, and whiskers are max to min. **** indicates p < 0.0001 in a non-parametric rank sum test. See also Figure S2 for qPCR validation of *CHD2* knockdown, cyclin E quantification, and spindle defect examples.

### An ASD patient-derived CHD2 missense variant prevents localization to the mitotic spindle

We next reasoned that studying patient derived ASD-associated missense mutations might provide additional insights into disease mechanisms beyond those observable in a knockdown model. In ASD exome sequencing studies (Satterstrom et al., 2020), 3 missense variants have been identified in *CHD2*. Of these, 2 have strong evidence for being pathological based on analysis of missense badness, PolyPhen-2, and constraint (MPC) scores (Samocha et al., 2017) (**Fig. 3A**). Therefore, we cloned and tagged these two variants, expressed them in *Xenopus*, and localized them in mitotic cells. Patient-derived missense variant CHD2^D856G^ did not localize to the mitotic spindle in *Xenopus*, and instead remained associated with DNA throughout mitosis (**Fig. 3B**). In contrast, patient-derived missense variant CHD2^G1174D^ localized to the mitotic spindle, similar to the reference sequence. Interestingly, CHD2^D856G^ is within the protein’s helicase domain, while CHD2^G1174D^ is within the DNA binding domain (**Fig. 3A**), suggesting that functional helicase activity, but potentially not DNA binding, is required for spindle re-localization during mitosis.

**Figure 3.**
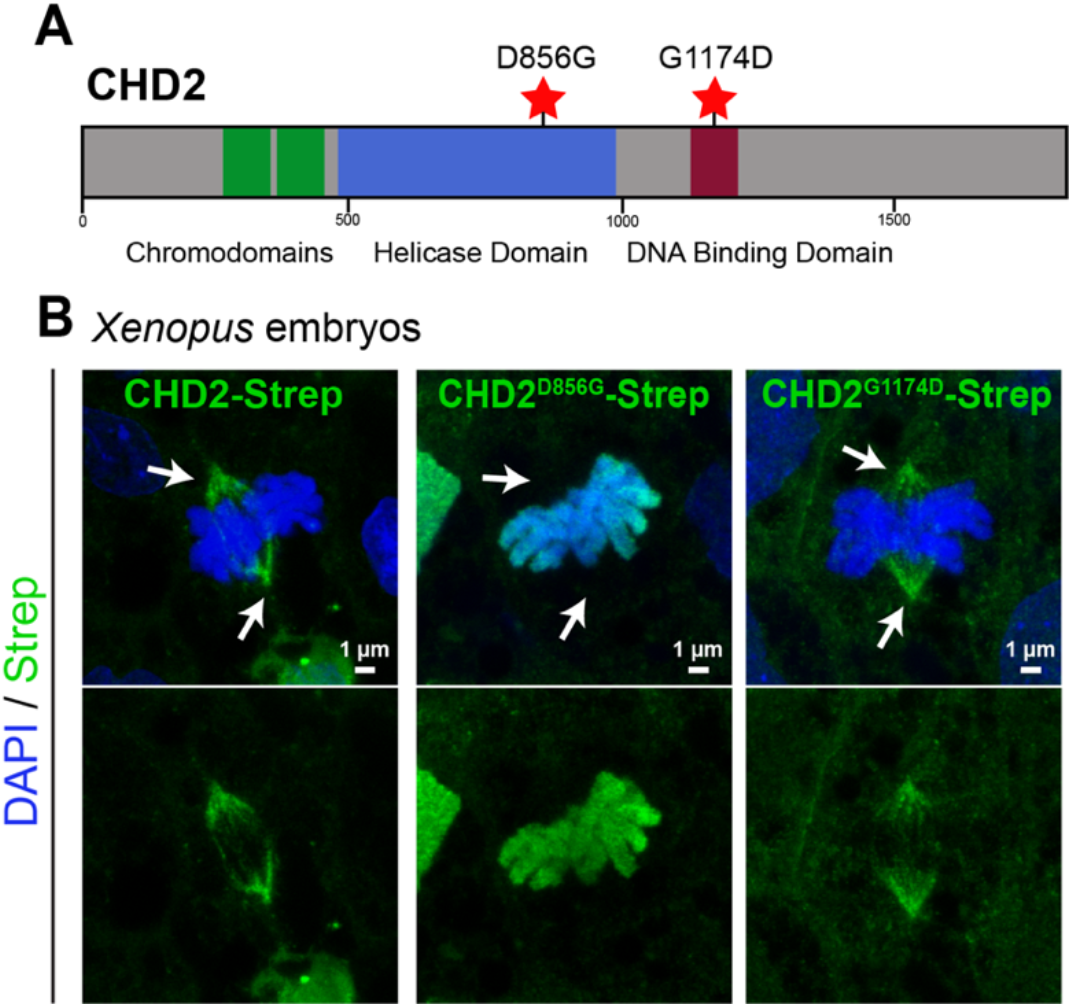
A CHD2 patient-derived missense variant disrupts spindle localization. (A) Schematic of Human CHD2 protein with functional domains and locations of likely pathogenic patient-derived missense variants annotated. Strep-tagged human CHD2 expressed in *Xenopus* localizes to mitotic spindles during mitosis, whereas the strep-tagged ASD patient-derived variant CHD2^D856G^ does not and instead remains localized to DNA (DAPI, blue). In contrast, ASD patient-derived variant CHD2^G1174D^ still localizes to mitotic spindles during mitosis.

### Microtubule-related proteins in general carry risk for autism

To test our second hypothesis–namely, that microtubule biology is related to ASD pathobiology in general–we queried a protein-protein interaction (PPI) network generated from affinity-purification mass spectrometry of centrosomal satellite proteins (Gheiratmand et al., 2019). Centrosomes are core microtubule-organizing centers, and the resulting PPI network contains many proteins involved in microtubule regulation (Gheiratmand et al., 2019). We first tested whether hcASD genes are overrepresented within this network. As this PPI network was generated in HEK293 cells, we restricted the list of hcASD risk proteins to those that have been previously shown to be readily detectable by quantitative mass spectrometry in these cells (Bekker-Jensen et al., 2017), generating a list of 81 hcASD risk proteins and a list of 623 centrosomal protein network interactors (**Table S2**). A significant number of ASD risk proteins (13 of 81 (16%)) overlapped between the hcASD risk gene list and the centrosomal satellite PPI (fold-enrichment 2.60, hypergeometric test p = 0.0013; **Fig. 4A**). Of these 13 overlapping proteins, 6 of them (*CHD8, SUV420H1/KMT5B, ARID1B, WAC, CREBBP, CTNNB1*) are annotated as “chromatin binding” (GO:0003682), an enrichment not expected by chance (odds ratio 11.95, Fisher’s exact test p = 0.00014).

**Figure 4.**
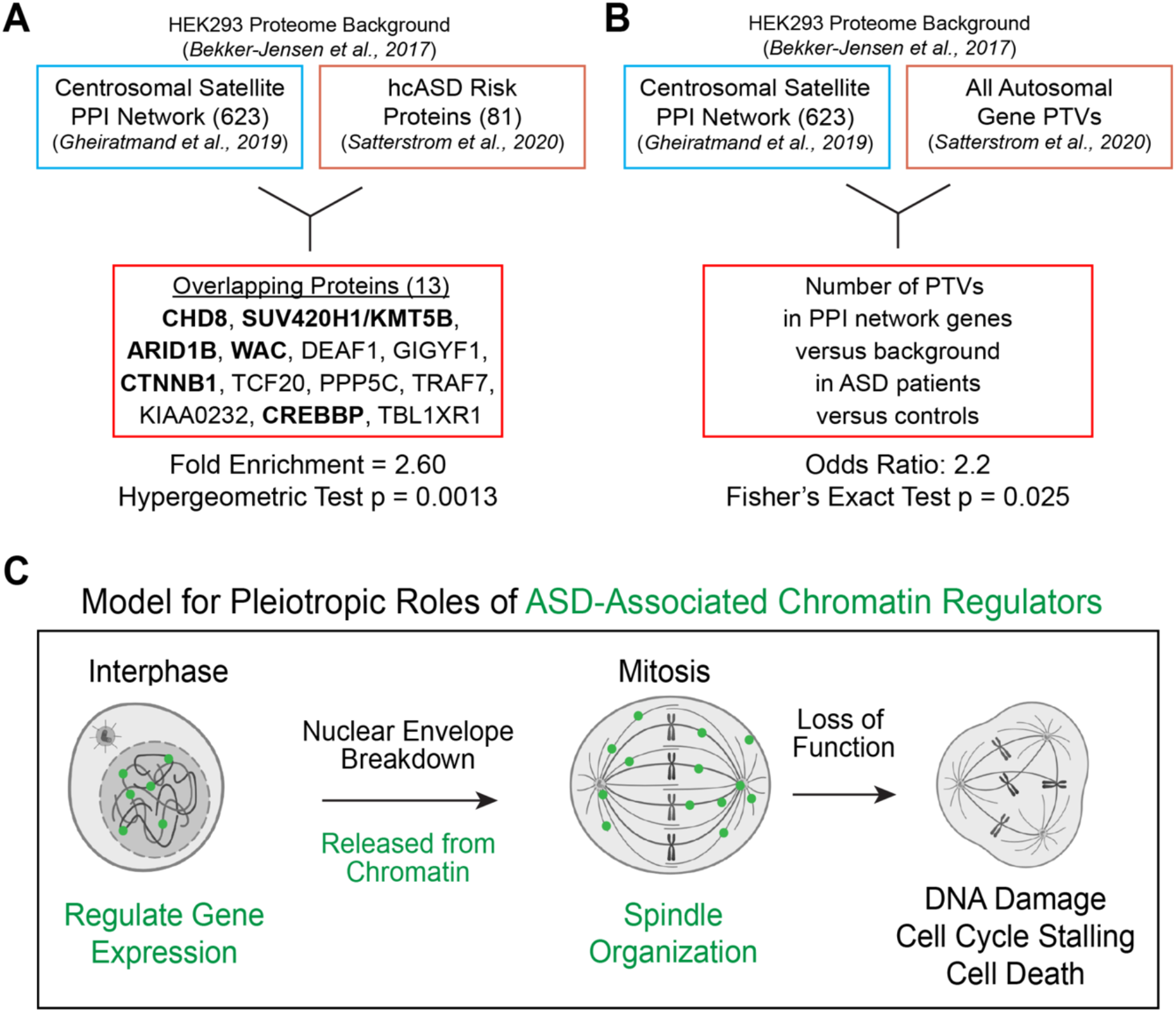
Enrichment of microtubule-related proteins in ASD. (A) High confidence ASD genes (Satterstrom et al., 2020) are overrepresented among microtubule-related proteins identified by affinity-purification mass spectrometry of centrosomal satellite proteins (Gheiratmand et al., 2019). A significant number of the overlapping genes (bold) encode proteins annotated as chromatin binding. (B) Genes encoding centrosomal proteins are more likely to carry protein truncating variants (PTVs) in ASD patients versus unaffected controls (Satterstrom et al., 2020). (C) Model for how ASD-associated chromatin regulators could play pleiotropic roles regulating chromatin and microtubules. During interphase, these proteins can localize to DNA, regulate chromatin, and control gene expression. However, after nuclear envelope breakdown during mitosis, these proteins relocalize to regulate microtubules and organize the mitotic spindle (modified from (Yokoyama, 2016)). In loss of function, microtubules are misorganized, the spindle does not form properly, and this leads to cell cycle progression defects, DNA damage, and ultimately cell death.

We next tested whether these centrosomal PPI network interactors were enriched within a large molecular interaction network constructed from high-confidence ASD risk proteins and the PCNet database of molecular interactions (Huang et al., 2018). Here we trimmed the PCNet-derived molecular interaction network to proteins present in the HEK293 background proteome, so as not to penalize the enrichment test for proteins that could not readily be detected in the centrosomal PPI network generation (Bekker-Jensen et al., 2017). A significant proportion of the centrosomal satellite PPI network interactors (**Table S2**) are present in the ASD interaction network (528 of 609 or 86.7%, fold-enrichment 1.2, hypergeometric test p = 6.6×10^−19^).

Finally, we asked whether presence in the centrosomal PPI network increases the odds that a gene carries a pathogenic mutation in an ASD patient. Indeed, we found that genes encoding proteins in this centrosomal PPI network are significantly more likely to contain protein truncating variants (PTVs) identified in ASD patients, as compared to non-network genes and unaffected control individuals (odds ratio 2.2, Fisher’s exact test p = 0.025; **Fig. 4B**).

## DISCUSSION

Multiple lines of evidence suggest an important role for tubulin biology in ASD pathogenesis. We previously identified a convergent role for ASD risk genes in cell proliferation and implicated microtubule regulation in this process for DYRK1A and KATNAL2 (Willsey et al., 2018, 2020, 2021). Others have described direct microtubule-related functions for ASD risk proteins including chromatin regulators (CHD3 and ADNP), a kinase (DYRK1A), and a microtubule-severing protein (SPAST) (Kuo et al., 2019; Ori-McKenney et al., 2016; Oz et al., 2014; Sillibourne et al., 2007). Here we have studied five genes selected based on their statistical evidence for association with ASD and their annotated function as chromatin regulators (ADNP, CHD8, CHD2, POGZ, SUV420H1/KMT5B), and observe that all five localize to microtubules *in vivo* and *in vitro*. We also describe a role for CHD2 in mitosis, with loss of function leading to mitotic spindle defects, cell cycle stalling, DNA damage, and cell death, all consistent with a role regulating microtubules. We therefore propose a model in which ASD-associated chromatin regulators perform pleiotropic roles depending on cell cycle phase (**Fig. 4C**). Specifically, during interphase, chromatin regulators localize to the nucleus and perform gene expression regulatory functions, but once the nuclear envelope breaks down and the cell enters mitosis, these proteins relocalize to organize the tubulin-based mitotic spindle, as has been proposed for other chromatin modifiers such as CHD4 (Yokoyama, 2016). Therefore, loss of function leads to mitotic spindle defects, cell cycle stalling, DNA damage, and cell death (**Fig. 4C**), which ultimately may intersect ASD pathobiology by causing defects in proliferation of progenitor cells.

This model and the results presented here offer potential insight into our prior findings showing that *ADNP, CHD8, CHD2*, or *POGZ* mutant *X. tropicalis* have an apparent expansion of proliferative neural progenitors, but ultimately a smaller brain size (Willsey et al., 2021). In consideration of our new work, our model now suggests that the apparent expansion of proliferative cells was most likely the result of cell cycle stalling during M phase and that the smaller brain size was likely due to subsequent cell death and/or perturbed proliferation. Consistent with this idea, patients with variants in *ADNP, CHD8, CHD2*, or *POGZ* commonly present with growth abnormalities and brain size changes (Assia Batzir et al., 2020; Bernier et al., 2014; Firth et al., 2009; Kawarai et al., 2018; Suls et al., 2013).

In general, the path from a growing list of large effect mutations to an actionable understanding of pathobiology has been complicated by, among other things, the extensive pleiotropy of ASD risk genes (Sestan and State, 2018; Willsey et al., 2021). As a case in point, from the earliest successes in rare variant gene discovery in ASD, genes annotated as chromatin regulators have been overrepresented among high confidence genes, leading to the conventional wisdom that pathobiology is due to dysregulation of gene expression (Valencia and Pașca, 2022). For example, the ASD-associated chromatin regulator CHD8 has been extensively studied with respect to its known role in chromatin biology. Interestingly however, ASD-associated loss-of-function mutations in *CHD8* appear to have a modest effect on global gene expression (Cotney et al., 2015; Sugathan et al., 2014), and transcriptional effects vary widely between studies (Wade et al., 2018), raising the possibility that CHD8 may play additional roles in ASD beyond regulating histones and gene expression. The work presented here broadens the likely relevant functions of such well-established chromatin regulators to include microtubule organization and underscores the value of including hypothesis-naive approaches to studying mechanistic convergence among a diverse set of risk genes as an alternative to relying on annotated function to identify pathobiology.

In addition to directly observing the impact of ASD associated mutations on microtubule biology, we provide strong evidence for the contribution of microtubule biology more broadly to ASD risk by leveraging *in silico* analyses of PPI resources. However, key questions remain regarding the relative contributions of transcriptional versus microtubule disregulation to ASD pathobiology. Specifically, for these “dual-function” genes it is unclear whether disruptions of chromatin, microtubules, or both underlie ASD pathobiology–especially because we cannot exclude the possibility that the microtubule-related defects seen in our CHD2 loss of function experiments are not due to the protein’s function at chromatin. Along the same line, we observe that a likely pathogenic patient-derived missense variant (CHD2^D856G^) does not localize to the mitotic spindle. However, a second likely pathogenic missense variant (CHD2^G1174D^) still localizes to the spindle. This could be for several reasons. First, this variant may localize correctly but aberrantly function at the mitotic spindle. Second, the missense mutation may not actually be pathogenic. Third, this variant may not impact microtubule biology. Additionally, it is unclear whether these missense variants also impact chromatin biology. Future work interrogating this and other patient-derived variants will be necessary to disentangle these possibilities. Given the precedent for chromatin regulators (including CHD3 and CHD4) to play independent and direct roles at both chromatin and microtubules, it is likely that these functions are separable, but such conclusions require a system where the functions can be tested in isolation, for example by leveraging the *Xenopus* cell-free cytoplasmic egg extract system (Gard and Kirschner, 1987; Heald et al., 1996). This will be important future work.

The tubulin code and associated post-translational modifications are rich and complex. There are 10 α-tubulin isotypes and 9 β-tubulin isotypes in humans, each with many different post-translational modifications and cell-type-specific expression patterns (Janke and Magiera, 2020). Deep complexity arises from the potential combinations of isotypes, modifications, and developmental context. The complexity of tubulin regulation underscores its diverse functions in development, ranging from mitosis to ciliogenesis to cell migration to synaptic transmission (Janke and Magiera, 2020; Lasser et al., 2018; Roll-Mecak, 2020). Understanding how ASD risk proteins intersect microtubules in all these contexts will be important, and determining which, if any, are relevant to pathobiology will be critical. Given the plethora of FDA-approved microtubule-targeting drugs from the field of oncology, identifying tubulin-related ASD mechanisms may offer a particularly tractable path for the development and testing of rational therapeutic strategies.

## MATERIALS AND METHODS

### Human ASD risk gene plasmid construction

ASD risk gene isoforms were selected based on gene expression in the developing cortex and the presence of exons with ASD patient variants using Clonotator software (www.willseylab.com/resources). Human cDNA was purchased from a third party company and cloned into pCDNA4 using In-Fusion cloning (Berrow et al., 2007) to add a dual-strep tag at either the N or C terminus (**Table S1**). Plasmid inserts were sequence-verified via Sanger sequencing. Patient-derived variants were created by site-directed mutagenesis using Q5 High Fidelity Master Mix (NEB Catalog #M0492L), T4 polynucleotide kinase (NEB Catalog# M0201L) and T4 DNA ligase (NEB Catalog# M0202L). Variants were selected by presence in Satterstrom et al., 2020 with an MPC score > 2 (Samocha et al., 2017).

### *Xenopus* care, husbandry, and microinjection

*Xenopus tropicalis* adult breeding animals originated in the Khokha lab (Yale, wild-type *Superman* strain), in the National *Xenopus* Resource (RRID:SCR_013731, (Pearl et al., 2012), wild-type *Superman* strain), or from Nasco (Fort Atkinson, WI, wild-type). Animals were cared for at the Willsey Lab aquatic facility in a recirculating system and used in accordance with an approved UCSF IACUC protocol (AN183565-02B). Wild-type animals of both sexes were used for these experiments. Natural matings and *in vitro* fertilizations were performed, where human chorionic gonadotropin (Sigma) was used for ovulation according to (Sive et al., 2007). Embryos were staged according to (Faber and Nieuwkoop, 2020). Clutch mates or internal within-animal tissue control cells were always used as controls. Parker Picospritzer III, Narishige micromanipulators, and Zeiss Stemi 508 microscopes were used to inject 2 nL of injection mix into one blastomere of two-cell stage *X. tropicalis* embryos. Plasmids were injected at 20 pg per blastomere. All injected reagents are detailed in **Table S1**. A fluorescent dextran (Thermo D34679) was always co-injected to label the injected side of the embryo, and injected animals were sorted according to fluorescence at neurula stages.

### *Xenopus* immunostaining, microscopy, and image analyses

Whole mount immunostaining of tailbud-stage animals was carried out according to (Willsey et al., 2018, 2020), including the bleaching step (this step was critical for permeabilization to visualize mitotic spindles in the epidermis), with dyes and DAPI (Thermo D3571) added during the secondary antibody incubation. All staining reagents are detailed in **Table S1**. Samples were mounted on glass slides in a vacuum grease well with PBS and coverslipped. Localization images were acquired on a Leica SP8 laser scanning confocal microscope with a 63x oil objective. Images were acquired as z-stacks at system-optimized intervals and processed in FIJI as maximum intensity projections. When fluorescence intensity was compared between samples, imaging acquisition settings were maintained between samples.

### HEK293 cell culture, transfection, staining, and imaging

Human HEK293 cells were maintained at 37°C in complete media consisting of DMEM with high glucose GlutaMAX Supplement (Gibco) and 10% Fetal Bovine Serum, and passaged at 20K cells/well using 0.25% Trypsin+EDTA (Gibco). Cells were transfected onto a 96-well optically-clear plate (Greiner CELLSTAR plates) that was coated overnight with 5μg/cm^2^ Fibronectin (R&D Systems) in PBS or onto glass coverslips in tissue culture plates. Transfections were performed using the PolyJet *in vitro* DNA transfection reagent kit (SignaGen Laboratories). 48 hours after transfection, cells were fixed in 4% paraformaldehyde at room temperature (RT) for 30 minutes. Cells were permeabilized for 15 minutes in PBS with 0.125% TritonX-100 (PBST), and blocked for 45 minutes with 2% bovine serum albumin in PBST (HEK Blocking Solution) at RT. Cells were incubated in primary antibodies diluted in HEK Blocking Solution overnight at 4°C, permeabilized for 15 minutes at RT, and incubated in secondary antibodies with DAPI (Thermo D3571) in HEK Blocking Solution for 1 hr at RT. Antibodies are detailed in **Table S1**. Cells were imaged on a Leica SP8 laser scanning confocal with 40x (for culture plates) or 63X (for glass slides) objectives. Images were acquired as z-stacks at system-optimized intervals and processed in FIJI as maximum intensity projections.

### Human iPSC-derived NPCs, immunofluorescence, imaging, and analyses

Non-targeting and *CHD2* CRISPRi iPSC-derived NPCs were generated as in Willsey et al., 2021. *CHD2* knockdown was confirmed by qPCR using primers 5’-CAGAGAGTGAGCCAGAACAAA-3’ and 5’-CCTTCCACTTGATGAGGTACTG-3’ (**Fig. S2**). NPCs were plated on Matrigel coated glass coverslips and fixed in 4% paraformaldehyde. Cells were then permeabilized for 10 minutes in PBS containing 0.2% Triton X-100 and blocked in PBS containing 0.1% Triton X-100 with 5% normal goat serum (NPC blocking solution) for 30 minutes at RT. Cells were incubated for 1hr at RT in primary antibodies (**Table S1**) diluted in NPC blocking solution, washed in PBT, incubated in secondary antibodies (**Table S1**) diluted in NPC Blocking Solution for 1 hr at RT, and washed in PBT and then PBS. Coverslips were mounted with DAPI (Thermo D3571) on glass slides before being imaged. Images were acquired using a Leica SP8 laser scanning confocal with 10x or 63x objectives. Images were acquired as z-stacks at system-optimized intervals, and processed into maximum intensity projections in FIJI.

The proportion of dividing cells (phospho-Histone H3 positive) with mitotic spindle defects was determined from images of the mitotic spindle (stained by β-Tubulin). All phases of mitosis (prometaphase, metaphase, anaphase, and telophase) were included. Spindles were considered normal if they were bipolar (two symmetric spindle poles) and abnormal if they were monopolar (one spindle pole), multipolar (more than two spindle poles), or disorganized (asymmetric bipolar) (see **Fig. S2B** for examples). Difference in spindle defect proportion was tested with a 2 × 2 contingency table and a chi squared test.

pH2AX puncta were quantified using the CellProfiler 4.1.3 Software (McQuin et al., 2018) as previously described in (Popova et al., 2021). Briefly, the “Speckle Counting” example pipeline was modified to produce robust identification of puncta. First, DAPI+ nuclei were identified using a diameter range of 70-250 pixels, threshold strategy “Adaptive”, and threshold method “Otsu”. The pH2AX channel was first enhanced and then masked, and puncta were subsequently counted using a diameter range of 7-10 pixels, threshold strategy “Adaptive”, and threshold method “Otsu”. A parent-child relationship was then assigned to group pH2AX puncta by DAPI+ nucleus, and the CellProfiler output was exported to Microsoft Excel. The number of pH2AX puncta per nucleus was plotted by condition and tested for statistical significance between conditions using an unpaired non-parametric rank sum test in GraphPad Prism 9. Cyclin B and Cyclin E mean fluorescence intensity (MFI) was quantified using the Analyze Particle function after thresholding in FIJI and tested for statistical significance between conditions using an unpaired non-parametric rank sum test in GraphPad Prism 9. For CCP3, cells with positive antibody staining were marked and counted manually in FIJI. Differences in the ratio CCP3 positive cells/total number DAPI+ cells between CRISPRi conditions were tested for statistical significance using an unpaired non-parametric rank sum test in GraphPad Prism 9.

### Systems biological analyses

We first created a custom background for high confidence ASD (hcASD) risk gene enrichment by intersecting the 11,215 proteins expressed in HEK293T cells (Bekker-Jensen et al., 2017) the 17,484 autosomal genes queried in (Satterstrom et al., 2020), resulting in 10,043 unique proteins. In this custom background, 81 out of the 102 hcASD genes (BFDR <= 0.1) (Satterstrom et al., 2020) and 623 out of the 662 unique centriolar satellite interactors from the centriolar satellites bait-prey protein interaction dataset (BFDR <= 0.01) (Gheiratmand et al., 2019) (**Table S2**). We identified 13 overlapping proteins and tested enrichment by hypergeometric test. We then assessed whether the 13 hcASD/centrosomal genes are enriched for genes that are annotated as “chromatin binding” in GO (GO:0003682). We defined the universe of genes to be the union of the 81 hcASD genes and the 623 centrosome genes (total of 691 genes, which includes 52 of 572 chromatin binding genes in GO:0003682). We conducted a Fisher’s exact test using a 2 × 2 contingency table using the variables hcASD + centrosome gene (yes/no) and chromatin binding (yes/no).

Next, we tested whether an interaction network built from PCNet (Parsimonious Composite Network) (Huang et al., 2018) around hcASD risk genes (Willsey et al., 2021) is enriched for these centrosomal protein interactors. To do this, we created a custom background by intersecting proteins from PCNet with all autosomal genes queried in ASD exome sequencing studies (Satterstrom et al., 2020) and proteins expressed in HEK293T cell lines (Bekker-Jensen et al., 2017), which resulted in 10,193 unique proteins. Using this background, 609 of the centrosomal interactors remained. 528 of these proteins were present in the hcASD network built from PCNet, and this enrichment was tested by hypergeometric test.

Finally, we performed a Fisher’s Exact Test to calculate the significance and odds ratio for de novo protein truncating variants (PTVs) (Satterstrom et al., 2020) occurring in centrosomal PPI genes (Gheiratmand et al., 2019) versus non-centrosomal but HEK293T expressed genes (Bekker-Jensen et al., 2017) in ASD patients versus unaffected control individuals.

## ACKNOWLEDGEMENTS

We thank Nolan Wong, Louie Ramos, and Will Figueroa for animal care; Juan Arbelaez and Milagritos Alva for lab maintenance; Tom Nowakowski for generous use of his confocal microscope; Kelsey Hennick for assistance quantifying pH2AX; Ashley Clement, Gigi Lopez, Sonia Lopez, and Linda Chow for administrative support. This work would not be possible without daily reference to the *Xenopus* community resource Xenbase (RRID:SCR_003280) and expertise and wild-type frogs from the National *Xenopus* Resource (RRID:SCR_013731).

## AUTHOR CONTRIBUTIONS

Conceptualization: HRW, AJW

Formal Analysis: ML, NS, KL, BW, HRW

Funding Acquisition: MWS, AJW, HRW

Investigation: ML, NS, YX, KL, SG, YZ, JD, KM, EB HRW

Methodology: ML, YX, JD, KM, HRW

Project Administration: HRW Resources: VD, SD

Supervision: HRW, AJW, MWS

Validation: ML, NS, SG, YZ, JD, HRW

Visualization: ML, NS, YX, SG, JD, HRW

Writing- original draft: HRW, AJW

Writing- review & editing: HRW, AJW, MWS

## DATA AVAILABILITY

Centrosomal satellite PPI data are publicly available at (Gheiratmand et al., 2019) and ASD risk gene data are publicly available at (Satterstrom et al., 2020).

## SUPPLEMENTAL FIGURES

**Figure S1.**
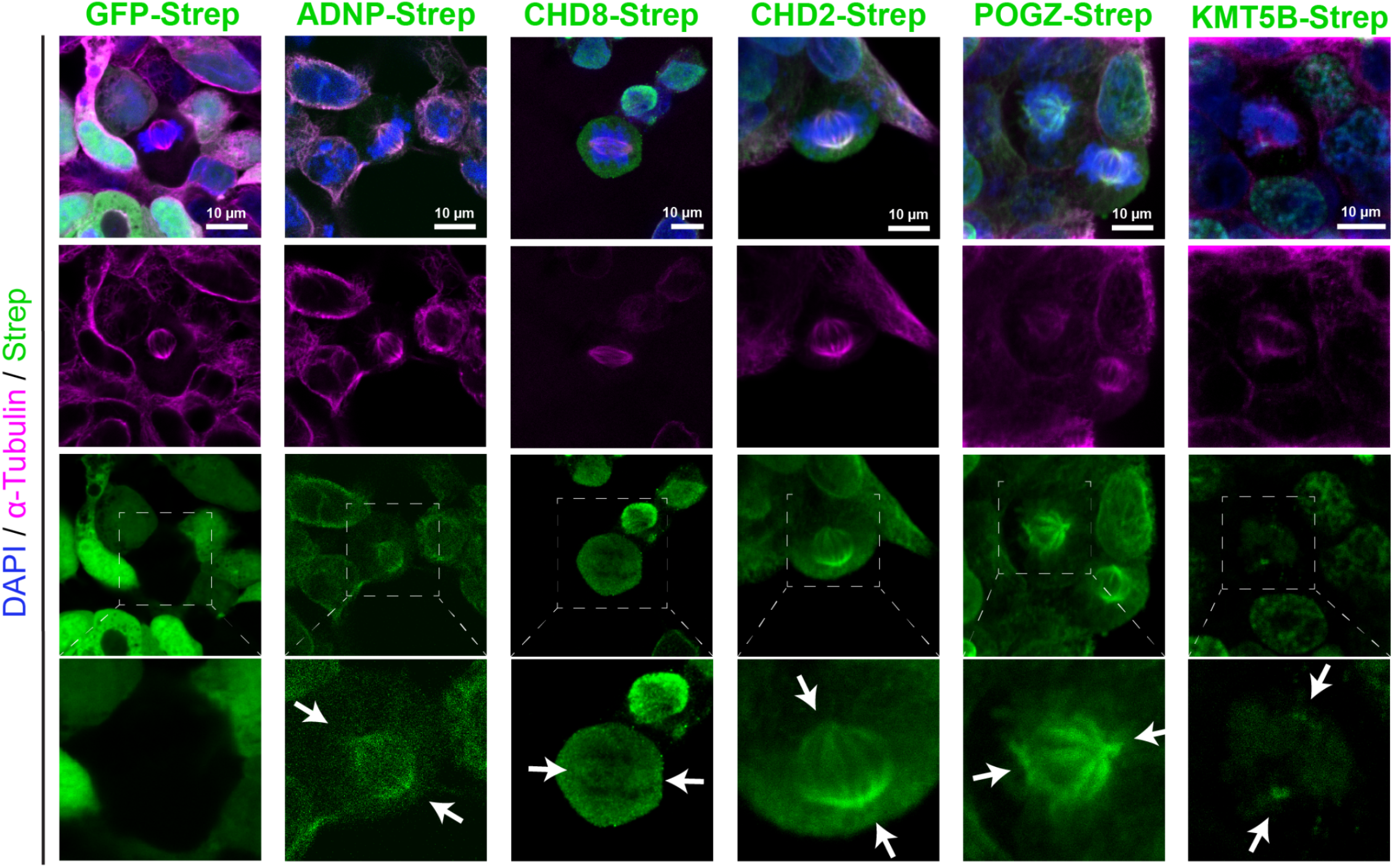
ASD-associated chromatin regulators localize to microtubules *in vitro*. ASD-associated chromatin regulators (ADNP, CHD8, CHD2, POGZ, and SUV420H1/KMT5B) localize to the mitotic spindle or centrosome when expressed in HEK293T cells, while a control GFP-strep construct does not. Bottom panel is a higher-magnification view of the boxed area in the panel above. Arrows point to the spindle poles. Related to Figure 1.

**Figure S2.**
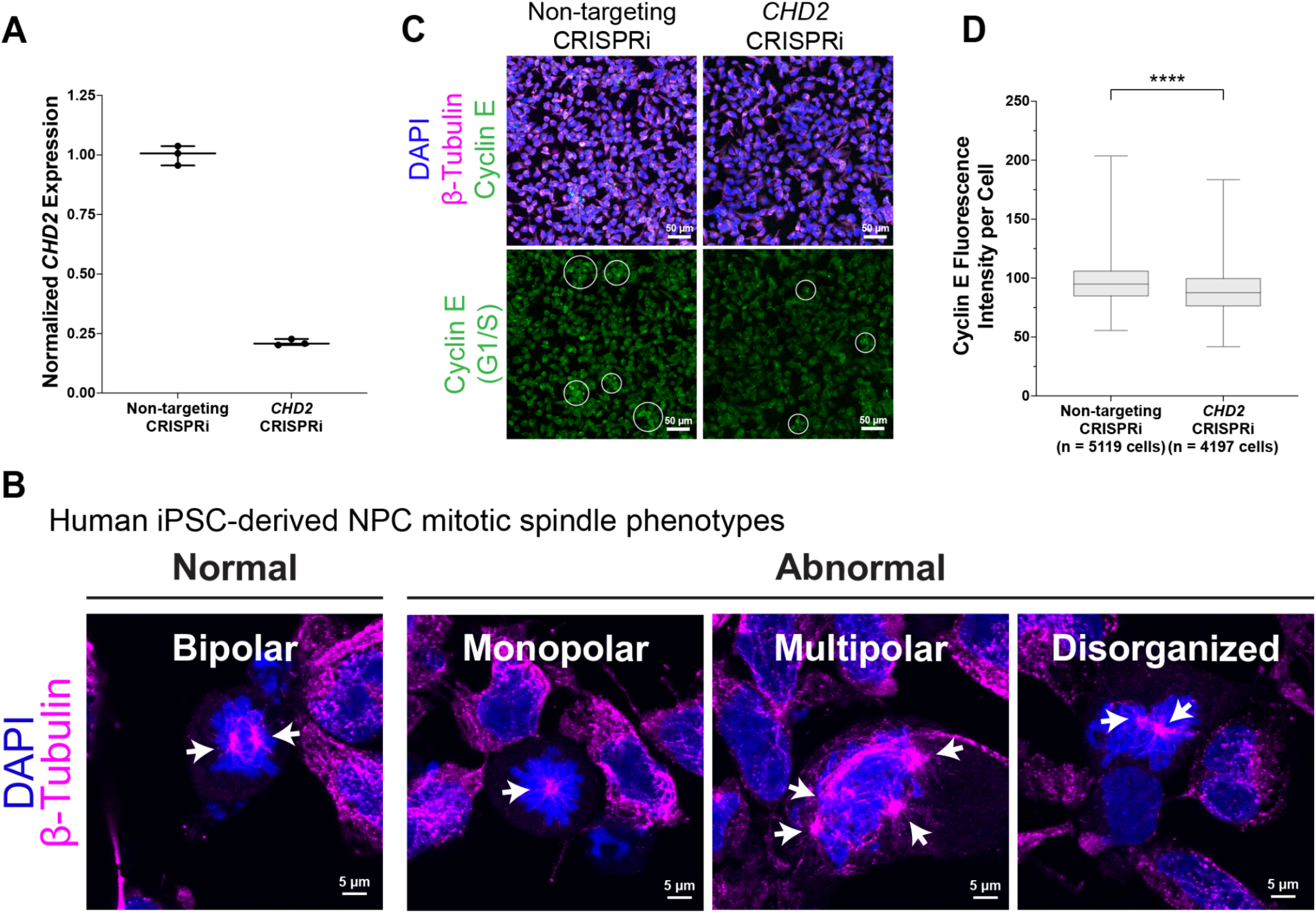
*CHD2* knockdown validation, spindle defect examples, and cyclin E staining. (A) *CHD2* expression is reduced following CRISPRi targeting of *CHD2*, compared to a non-targeting control CRISPRi line, as determined by qPCR. (B) Examples of normal and abnormal mitotic spindles for Figure 2A. (C) CRISPRi of *CHD2* causes a significant decrease in Cyclin E (G2/M marker) fluorescence per cell compared to a non-targeting CRISPRi line. (D) Quantification of C. Box is 25-75% interquartile range, line is median, and whiskers are max to min. **** represents p < 0.0001 by rank sum test. Related to Figure 2.

